# Response of sugarcane rhizosphere soil fungal communities on a temporal gradient to changes in critical growth periods

**DOI:** 10.1101/2022.06.21.497116

**Authors:** Zhaonian Yuan, Qiang Liu, Ziqin Pang, Yueming Liu, Fallah Nyumah, Chaohua Hu, Wenxiong Lin

## Abstract

Understanding the normal variation of the sugarcane rhizosphere fungal microbiota throughout its life cycle is essential for the development of agricultural practices for fungal diseases (e.g., sugarcane tip rot, sugarcane red rot, and sugarcane smut) and ecological health associated with the microbiota. Therefore, we performed high-throughput sequencing of 18S rDNA of soil samples using the Illumina sequencing platform for correlation analysis of sugarcane rhizosphere microbiota time series, covering information from 84 samples in four growth periods. The results revealed that the sugarcane rhizosphere fungi possessed the maximum fungal richness in July (Tillering). Rhizosphere fungi are closely associated with sugarcane growth, including Ascomycota, Basidiomycota, and Ochrophyta showed high abundance in a stage-specific manner. Through the Manhattan plots, 11 fungal genera were found to show a decreasing trend throughout the sugarcane growth period, and two fungal genera were significantly enriched at three stages of sugarcane growth (*p* < 0.05) including *Pseudallescheria* and *Nectriaceae*. In addition, soil pH, soil temperature (Tem), total nitrogen (TN) and total potassium (TP) were important drivers of fungal community structure at different stages of sugarcane growth. Using structural equation modeling (SEM) found that sugarcane disease status showed a significant and strong negative effect with selected soil properties, suggesting that poor soil may increase the likelihood of sugarcane disease. In addition changes in sugarcane rhizosphere community structure over time were mainly influenced by stochastic factors, but the contribution decreased to the lowest value after the sugarcane root adaptation system was stabilized (Maturity).

**IMPORTANCE:** Rhizosphere microbes are closely related to plant growth, and more studies have shown that the rhizosphere fungal microbial community has an important influence on plant health and growth status. However, little is known about the response of the rhizosphere fungal community to plant growth during the critical plant reproductive period. In this study, we analyzed the important response of the rhizosphere fungal community of sugarcane through the pattern of abundance changes in its critical growth nodes by various methods to investigate the subtle changes in the assembly of the rhizosphere fungal community with the growth of sugarcane. Our work provides innovative ideas for the prevention of soil-borne diseases in plants and also provides a solid basis for the development of microbial models of crops rhizosphere soil.

## INTRODUCTION

The microbial community in the rhizosphere environment is critical for the health of land plants and sustainable soil development (1). As more attention is paid to the relationship between plants and soil microbiome, comprising interacting microbial and plant root systems and genetic components (2-4), it is apparent that much more exploration is needed of what constitutes normal temporal variations in rhizosphere microbial community structures and compositions. Variations in the rhizosphere microbiome and between different plants and life-cycle stages have a lot of space for exploration. As we all know, after germinating in the soil, plant roots begin to recruit microbial communities closely related to their growth through the accumulation of root secretions, rhizosphere deposition and nutrient uptake. Thus, the growth activities of plants during different periods may cause changes in rhizosphere microbial communities and functions. For instance, model plants (*Arabidopsis thaliana* and *Medicago truncatula*) can retain certain fungal populations in the soil through certain chemicals secreted by the root system(5). In addition, advances in sequencing technology provide a foundation for the research on the dynamic changes of microbes in the rhizosphere of plants in space and time (6). Similarly, these types of studies are needed to understand the immigration and emigration patterns of microorganims between different parts of the plant, between different points of time and between different plants and soil environments (7-9), especially the related research of time series. For example, Zhang et al.,(2018) by a machine learning approach, they identified biomarker taxa and established a model to correlate root microbiota with rice resident time in the field (10), and Li et al., (2020) analyzed the variation of the fungal community structure in the rhizosphere of Gardenia, which provided a new theoretical for further study on the rhizosphere mechanism (11). In addition, as an important economic crop of food and biofuel, sugarcane is constantly expanding its planting the area under the condition of global warming (12,13). The study on the time change of the sugarcane rhizosphere microbiome has great significance in better understanding the interaction between sugarcane and rhizosphere microbiota (14). For example, sugarcane growth is facing a huge threat due to Yellow Canopy Syndrome (YCS) in Australia. YCS is a kind of largely undiagnosed plant disease impacting sugarcane growth across Queensland, Australia, causing huge yield losses(15). Studying the time changes of sugarcane rhizosphere microorganisms can provide a new way of thinking about the solution of YCS. Besides, given the sugarcane growing importance (16,17), identifying ways to the healthy growth of sugarcane and increase productivity in a sustainable manner is critical and, hence, the potential to harness microorganisms from the sugarcane rhizosphere has recently gained more attention (18-20). For instance, rhizosphere microorganisms are involved in the solubilization of potassium-containing minerals (21). And sugarcane smut disease caused by a fungus *Sporisorium* scitamineum is a limiting factor to cane production. It is threatening the sugar industry. Juma and Musyimi’s research showed that the selected isolates from sugarcane rhizosphere microflora had evidently antagonistic activity against the *Sporisorium* scitamineum hence recommended as a potential biocontrol agent for this pathogen (22). Another example, She et al., (2021) found maize rhizosphere microorganisms are critical for facilitating microbiome bioremediation for soil affected by neutral-alkaline mining (23). Moreover, understanding how the sugarcane rhizosphere microbiome changes with seasons would provide important data and could in the long-term result in the identification of potential management strategies for healthy cultivation of sugarcane, including an improved breeding process taking the microbe into account, or better targeted biological prevention and treatments (24). Here, we used sugarcane as a research object to fill critical knowledge gaps that will accelerate our exploration process to successfully harness rhizosphere microbiomes to sustainably enhance a sugarcane productivity. This also has reference significance for other crops in nature. Our study had three main objectives: (i) to determine the drivers of the sugarcane fungal community under field conditions; (ii) to understand the seasonal trends of fungi in the rhizosphere of sugarcane; and (iii) to explore fungi species closely related to changes in sugarcane growth period. To achieve this, we examined fungal community assemblages of sugarcane rhizosphere soil through Illumina amplicon sequencing of samples collected from young to mature sugarcane (May, July, September, November). Our results provide more foundation into the process of sugarcane root microbiota exploration.

## MATERIALS AND METHODS

### Sugarcane planting

In the spring of 2017, sugarcane cultivar, FN 41, was grown in a separate sugarcane experimental field in China to track the rhizosphere microbiota change procedure during the entire sugarcane growth cycle. To avoid seed-canes surface-associated microbes, before planting, the sugarcane seed-canes were used 0.1% carbendazim solution to soak for 10 minutes. After soaking, soaked the seed-canes with water for 24 hours (24), then sugarcane seed-canes after soaking were transferred to fields on Baisha experimental station (119° 06’ E, 26° 23’ N). A randomized group design was used, with an area of 1,008 m^2^, 8 rows per zone, 5 m row length, 1.2 m row spacing, 48 m^2^ per plot, and 3 replications, distributed in randomized groups in each plot. The planting height of sugarcane was 1/2 of the row height. To measure the stalk height and stem diameter of sugarcane in each period, twenty-one sugarcane were randomly selected in each field and measured with a measuring tape and vernier caliper. Meanwhile, the number of effective plants, diseased plants and total plants were also counted.

### Sample collection

The experiment started on the 3rd of January 2017. The soil samples were collected at the Baisha Experimental Station of the National Sugarcane Research Center of Fujian Agriculture and Forestry University (subtropical monsoon climate, the annual average temperature 19.5°C, the annual average precipitation is 1,673.9 mm). Using the “S-shaped sampling method” to select 21 sampling points in the sugarcane field (26). According to the shaking-off method of Riley & Barber (27), the soil adhered to the sugarcane roots was brushed with a small sterile brush, and soil samples were finally collected, sieved at 2.0 mm, and stored in a refrigerator at -20 °C (28). The samples were collected from the sugarcane seeding stage (May 4, 2017), tillering period (July 2, 2017), elongation period (September 17, 2017), and maturity period (November 29, 2017), with a total of 84 samples (4 periods, 21 repetitions).

### Determination of pokkah boeng disease of sugarcane and soil physio-chemical properties

Soil suspension with water (1:2.5 WV^−1^) was prepared to estimate soil pH using a pH meter (PHS-3C, INESA Scientific Instrument Co., Ltd, Shanghai, China) (29). The soil temperature (Tem) was measured with the Soil Temperature Detector (Model: JC-TW, China). The available nitrogen (AN) was measured by the alkaline hydrolyzable diffusion method (30), Soil total N (TN) and organic matter (OM) were determined by Kjeldahl digestion and determined by oil bath–K_2_CrO_7_ titration method (31, 32). C:N is the ratio of soil total N to organic matter. Soil total K (TK) and total P (TP) were determined by digestion with HF-HClO_4_, followed by flame photometry and molybdenum-blue colorimetry, respectively (33,34). Available K (AK) was extracted by ammonium acetate and determined by flame photometry (35). Available P (AP) was extracted by sodium bicarbonate and determined using the molybdenum blue method (36). Field judgment of pokkah boeng disease of sugarcane was mainly based on the symptoms described in previous studies (37) and divided into three types: chlorotic phase, acute phase or top-rot phase and knife-cut phase (associate with top rot phase).

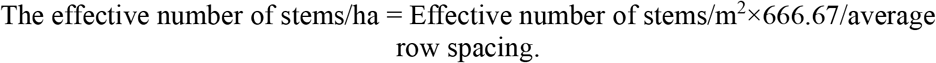

### DNA extraction and PCR amplification

For each soil sample, soil DNA was extracted using a Power Soil DNA Isolation Kit (MoBio Laboratories Inc., Carlsbad, USA) according to the manufacturer’s instructions. A NanoDrop 2000 spectrophotometer (Thermo Scientific, Waltham, MA, USA) was employed to assess the concentration and quality of soil DNA, was used. Amplification of 18S rDNA gene fragments was carried out using primers set SSU0817F/SSU1196R (38). The reaction conditions used for DNA amplification were: 95 °C for 3 min, followed by 35 cycles of 95 °C for 30 s, 55 °C for 30 s, and 72 °C for 45 s, with a final extension at 72 °C for 10 min (GeneAmp 9700, ABI, California CA, USA). PCR reactions were carried out in triplicate in a 20 μL mixture containing 2 μL of 2.5 mM sNTPs, 4 μL of 5× Fast Pfu buffer, 0.4 μL of Fast Pfu polymerase, 0.4 μL of each primer (5 μM), and template DNA (10 ng). Extraction of amplicons was carried out using an AxyPrep DNA Gel Extraction Kit (Axygen Biosciences, Union City, CA, USA). Later, the DNA was quantified using QuantiFluor™-ST (Promega, Madison, WI, USA). Purified amplicons were pooled in equimolar and paired-end sequenced (2 × 250) on an Illumina MiSeq platform (Majorbio, Shanghai) following to the standard procedures.

### Sequences and statistical analysis

The UPARSE standard pipeline was used to analyze the sequence data (39). Briefly, sequences with short reads (< 250 bp) were filtered out before to downstream analysis. Sequences with ≥ 97% similarity were clustered into OTUs. All sequences were assigned using the RDP classifier to identify taxa with a confidence threshold of 0.8 (40). We selected these OTU with 97% similarity, then used Mothur to calculate the Alpha diversity index under different random sampling (41), then used R to draw the rarefaction curves. The DPS was used to analyse the variance of the soil physical and chemical properties, and the significance was calculated based on the Tukey test (*p* < 0.05) (42). RDA was used to visualise the relationship between fungal communities and soil environmental factors. Network analysis used R to calculate the correlation between the factors (Spearman correlation), and Cytoscape (version 3.6.1) was used to adjust and visualise the results (43). Analysis of differential OTU abundance and taxa was performed using a DESeq2 of the R package, and then we used a Manhattan plot to visualize the results (R 3.6.0). Structural Equation Modeling (SEM) was performed using IBM SPSS Amos 26. Maximum likelihood estimation with standard errors was also used (44). The nearest-taxon-index (NTI) and βNTI (999 random) were used to quantify changes in rhizosphere fungal phylogeny over time, and the two indices were calculated using the package “picante” (45,46). The heatmap and functional annotation and maximum likelihood trees were done using the majorbio platform (http://cloud.majorbio.com). FUNGuild classification map was completed using STEMP (version 2.1.1), comparing the two seasons, a statistical test was Welch’s t-test, 95% confidence intervals, *p* < 0.05. Network diagramming and parameter calculation using R, Cytoscape and UCINET 6 together (47).

## RESULTS

### Seasonal changes of soil properties and phenotypic indicators of sugarcane

The results showed that the height and stem diameter of sugarcane plants increased continuously over time, with the rate of diseased plants reaching the highest in September (Table 1). We also investigated the temporal variation of soil biochemical properties collected in May, July, September and November during the different growth stages of sugarcane. Our analysis showed that soil biochemical properties varied with time. Soil properties such as soil pH and AN decreased considerably (*p* < 0.05), with September and November recording the lowest soil pH, while May and recorded the highest, followed by the month of July. In addition, the soil OM and TP were significantly higher in the month of September compared to May, July, November (*p* < 0.05). We also observed that May recorded the significant amount of soil TN relative to the other months. Compared to September, considerable amount of soil AP was recorded compared to the other months. In comparison to July and November, soil TK revealed significant improvement (*p* < 0.05) in May, followed by September. However, AK soil showed no significant difference among all the different time point. Soil temperature significantly peaked (*p* < 0.05) in July and September relative to May and November (Table 2).

**Table 1.** Changes in phenotypic indicators of sugarcane at different fertility stages

|  | Plant height(cm) | Stem diameter(mm) | Total number/ ha | Valid number/ ha | Effective ratio(%) | Disease ratio(%) |
| --- | --- | --- | --- | --- | --- | --- |
| May | 42.49±1.37<sup>c</sup> | 14.70±0.80<sup>c</sup> | 3270 | 3185 | 97.40 | 1.4 |
| July | 56.30±2.10<sup>c</sup> | 23.70±1.49<sup>b</sup> | 3473 | 3307 | 95.22 | 5.1 |
| Sep | 122.90±4.54<sup>b</sup> | 27.90±1.25<sup>b</sup> | 2940 | 2103 | 71.52 | 19.8 |
| Nov | 219.67±5.98<sup>a</sup> | 36.67±1.01<sup>a</sup> | 2950 | 2528 | 85.70 | 15.1 |

Note: Different letters in each column indicate significant difference among the treatments at $p < 0.05$ .

**Table 2.** Changes of soil nutrients in sugarcane at different growth stages

|  | May | July | Sep | Nov |
| --- | --- | --- | --- | --- |
| pH | 6.22±0.04<sup>a</sup> | 5.83±0.05<sup>b</sup> | 5.55±0.06<sup>c</sup> | 5.38±0.09<sup>c</sup> |
| OM/(g kg<sup>-1</sup>) | 26.78±0.89<sup>b</sup> | 25.03±0.54<sup>b</sup> | 30.80±1.91<sup>a</sup> | 26.71±1.19<sup>b</sup> |
| TN/(g kg<sup>-1</sup>) | 2.74±0.04<sup>a</sup> | 2.45±0.02<sup>b</sup> | 2.39±0.03<sup>b</sup> | 2.40±0.02<sup>b</sup> |
| TP/(g kg<sup>-1</sup>) | 0.80±0.01<sup>b</sup> | 0.78±0.01<sup>b</sup> | 0.92±0.02<sup>a</sup> | 0.82±0.01<sup>b</sup> |
| TK/(g kg<sup>-1</sup>) | 23.25±0.20<sup>a</sup> | 16.58±0.44<sup>c</sup> | 19.65±0.24<sup>b</sup> | 17.18±0.28<sup>c</sup> |
| AN/(mg kg<sup>-1</sup>) | 182.06±5.73<sup>a</sup> | 168.75±3.12<sup>b</sup> | 127.01±1.67<sup>c</sup> | 125.04±3.46<sup>c</sup> |
| AP/(mg kg<sup>-1</sup>) | 78.61±1.87<sup>ab</sup> | 82.08±1.19<sup>a</sup> | 65.47±1.48<sup>c</sup> | 75.46±2.39<sup>b</sup> |
| AK/(mg kg<sup>-1</sup>) | 155.99±4.08<sup>a</sup> | 149.97±3.49<sup>a</sup> | 136.11±5.70<sup>ab</sup> | 125.11±14.72<sup>b</sup> |
| C:N | 9.84±0.35<sup>b</sup> | 10.23±0.27<sup>b</sup> | 12.89±0.80<sup>a</sup> | 11.15±0.52<sup>ab</sup> |
| Tem | 24.49±0.04<sup>b</sup> | 30.36±0.06<sup>a</sup> | 30.22±0.04<sup>a</sup> | 19.37±0.03<sup>c</sup> |

Note: temporal variation of soil biochemical properties during sugarcane growth stages. Different letters in each column indicate significant difference among the treatments at $p < 0.05$ .

### Season variation of fungal diversity

The total number of sequences in the raw sequencing data is 4,592,374 with an average length of 401.21, the shortest sequence length is 221 and the longest sequence length is 453. For details, see Annexes-1. Fungal diversity (Shannon and Simpson), richness (ACE) and Sobs (number of observed species) were assessed during the different time points. It was observed that fungal diversity increased substantially (*p* < 0.05) in July compared with the other months (Table 3 and Fig. S1A). Moreover, compared to November, fungal richness and Sobs were significantly enriched (*p* < 0.05) in July, followed by May and September (Table 3, Fig. S1C and D). In addition, the dilution curve showed a sufficient amount of sequencing data (Fig. S2).

**Table 3.** Alpha diversity indices for different periods

|  | May | July | Sep | Nov |
| --- | --- | --- | --- | --- |
| Sobs | 310.81±6.34<sup>b</sup> | 388.48±6.79<sup>a</sup> | 328.62±6.46<sup>b</sup> | 277.67±12.15<sup>c</sup> |
| ACE | 396.43±9.88<sup>b</sup> | 466.04±9.49<sup>a</sup> | 415.22±9.08<sup>b</sup> | 338.92±14.47<sup>c</sup> |
| Shannon | 3.36±0.08<sup>b</sup> | 3.83±0.04<sup>a</sup> | 3.52±0.06<sup>b</sup> | 3.35±0.05<sup>b</sup> |
| Simpson | 0.1±0.02<sup>a</sup> | 0.05±0.003<sup>b</sup> | 0.08±0.01<sup>ab</sup> | 0.09±0.01<sup>a</sup> |

Note: temporal variation of α-diversity during sugarcane growth stages. Different letters in each column indicate significant difference among the treatments at $p < 0.05$ .

We carried out Principal Coordinates Analysis (PCoA) to explore and visualise similarities or dissimilarities in fungal community composition in all the samples collected during the different growth stages of sugarcane. Experimental design map of rhizosphere fungi microflora changes of FN 41 sugarcane during the whole life cycle (Fig. 1A). The analysis revealed that fungal community composition was clustered together in the different samples, and the soil samples shifted with different seasons in the first axis (Fig. 1B).

**Fig. 1.**
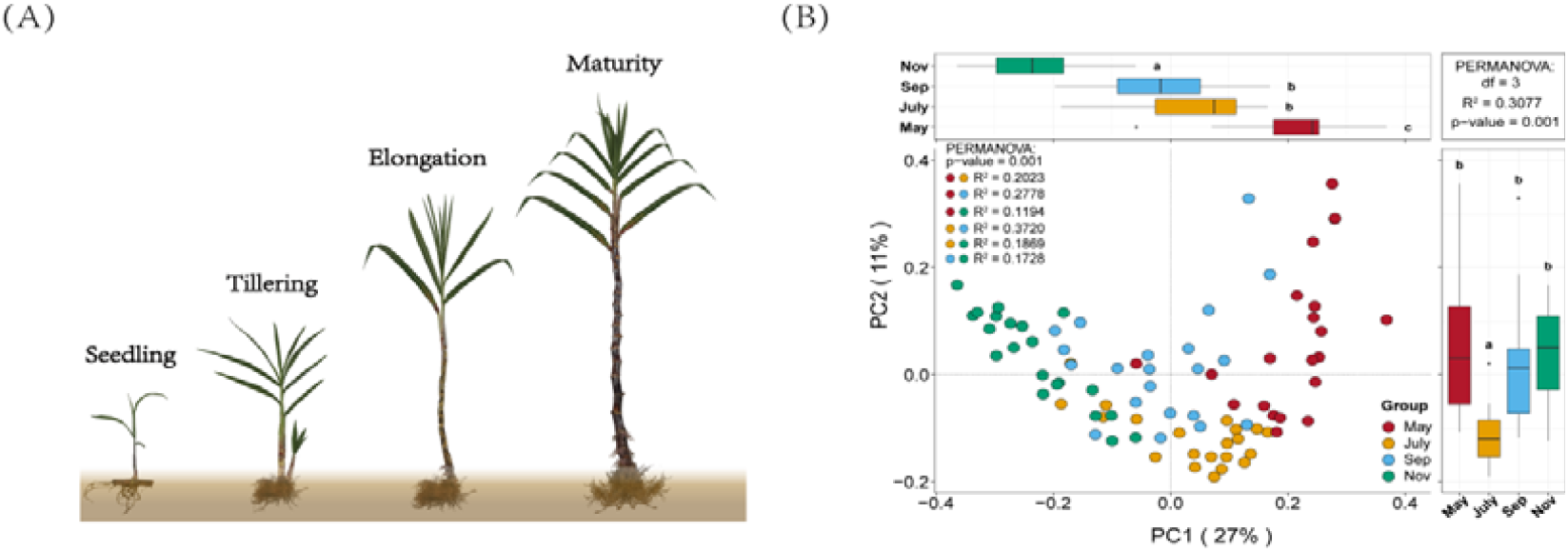
Fungal community composition under different sugarcane growth stages (A), as displayed by principal coordinates analysis (B). Each growth stage in is represented by: May (seedling stage), July (tillering stage), September (elongation) and November (maturity), all p-values were 0.001, and different colors represented different seasonal groups.

### Correlation between fungal alpha diversity and soil biochemical properties

The relationship between soil environmental variables and fungal alpha diversity was investigated using Pearson’s correlation coefficients (Fig. 2A). The analysis demonstrated that soil temperature (Tem) exhibited a strong and positive association with fungal Sobs and ACE, Shannon and PC1. However, soil Tem was negatively associated with PC2. Furthermore, soil pH revealed a significant positive correlation with PC1, Sobs and ACE. Soil TN and AN were significantly and positively associated with PC1. Soil TN also exhibited a positive relationship with PC2, while TK had a strong positive association with PC1 and PC2. We also noticed that soil TP demonstrated a positive correlation with PC2. On the other hand soil TN and TK were negatively related to Shannon. Meanwhile, regressions analysis was conducted to further confirm the association among soil environmental variables, fungal ACE (richness), Shannon (diversity), PC1 and PC2 during the different time points (Fig. S3). The analysis demonstrated that soil pH showed a positive relationship with fungal richness, soil AN was demonstrated a positive association with fungal richness as well. Moreover, soil Tem exhibited a positive relationship with fungal richness particularly in July and September (Fig. S3A). Soil pH and C/N showed a positive correlation with fungal richness in July, whereas AN showed a strong positive correlation with fungal richness in May. In July and September, soil Tem revealed a strong positive relationship with fungal richness (Fig. S3B). In May, soil pH, TN, AP, AN and AK were positive relationship with PC1, while in July and September, soil Tem was positively related with PC1 (Fig. S3C). Whereas in May, TN and TK had a positive association with PC2, while in September and November soil TP and AK demonstrated a positive correlation to PC2, respectively (Fig. S3D).

**Fig. 2.**
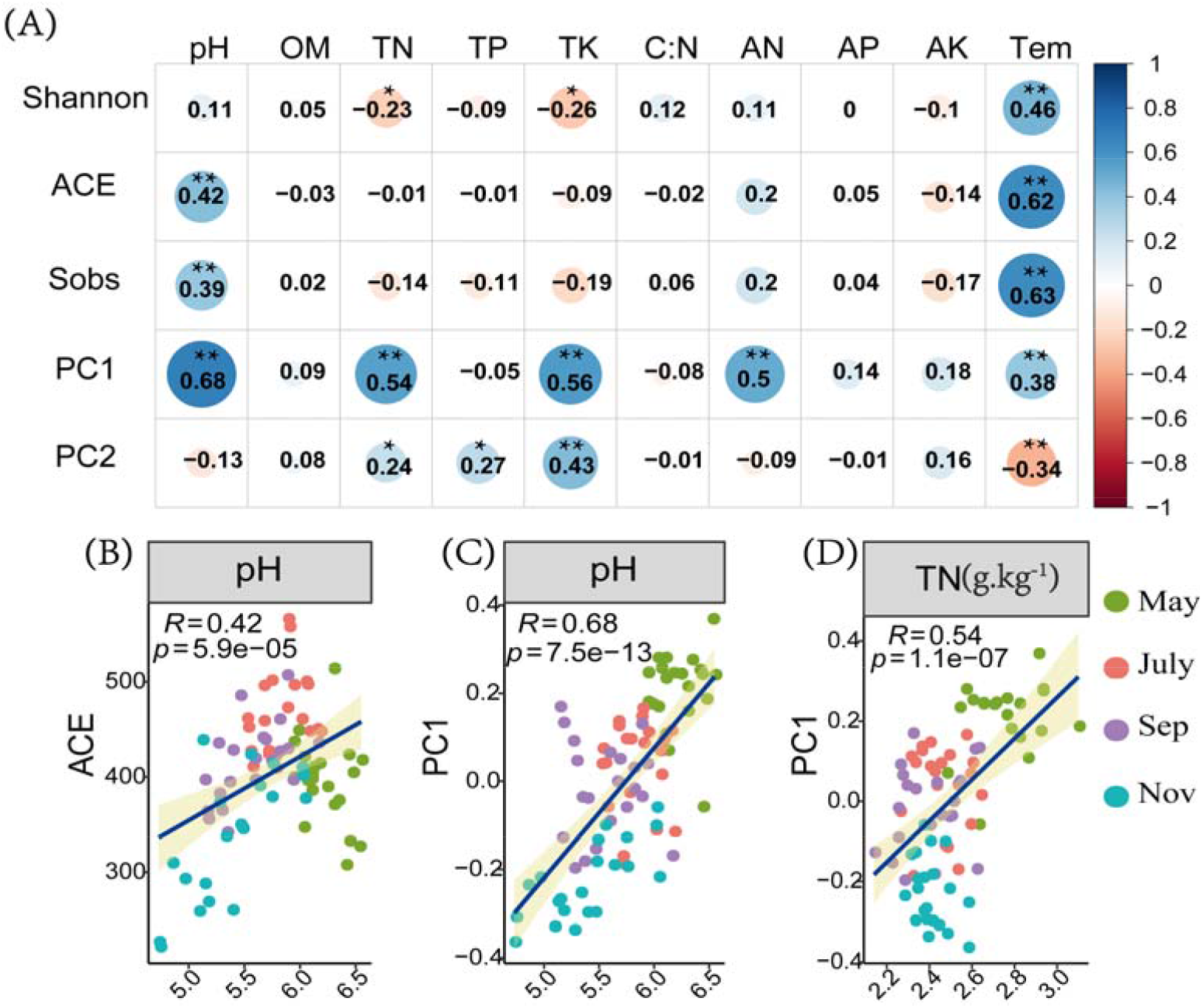
Pearson’s correlation coefficients for soil physio-chemical properties and fungal alpha diversity, PC1 and PC2 (A), and regressions analysis among soil environmental variables, fungal ACE (richness) and PC1 and PC2 sampled during the month of May, July, September and November (B). * *p* < 0.05, ** *p* < 0.01, *** *p* < 0.001.

### Relative abundance of fungal community composition

Fungal relative abundance during the different months was assessed at the phylum level. The analysis showed that Ascomycota, Basidiomycota, Chytridiomycota, Glomeromycota and Ochrophyta were dominant in the entire sample. However, Chytridiomycota, Ochrophyta, Platyhelminthes and Nucleariida were significantly more (*p* < 0.05) in July compared to the other months (Fig. S4A, S4B, S4F and S4I). Additionally, Ciliophora and Choanoflagellida decreased significantly in May relative to the other months, while Basidiomycota increased significantly (*p* < 0.05) in July and May compared to other treatments (Fig. S4C, S4D and S4H). Our analysis also showed that Glomeromycota reduced considerably in September relative to July and November, but the Eukaryota trend did not change during the entire period (Fig. S3E and S3G). Venn diagram analysis demonstrated that 25, 53, 20 and 15 unique fungal OTUs were detected in May, July, September and November, respectively (Fig. 3B). Moreover, among the soil samples collected in the various months, 511 enriched OTUs were shared among the different time points. In July, the highest number of enriched OTUs were recorded compared with the other sampling time.

**Fig. 3.**
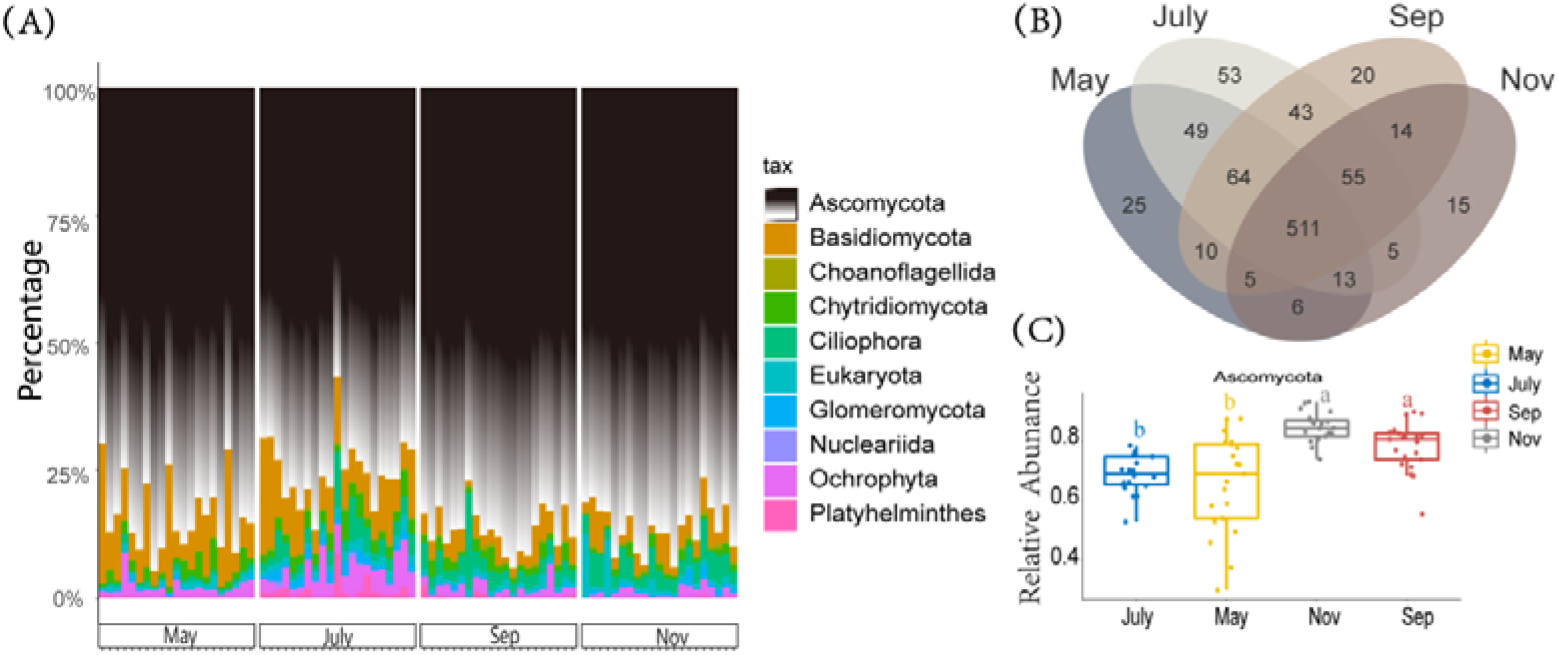
Different stages influences the diversity profiles of the rhizosphere soil microbiota. (A) Stacked graphs showed that relative abundances of the main phyla according four different growth periods, it was from twenty one soil samples of each period. (B) A Venn diagram of operational taxonomic units (OTUs) among four seasons (May, July, September, November). (C) The box-plot of relative abundance of *Ascomycota*, the rhizosphere fungus of sugarcane in different seasons(*p* < 0.05).

### Relationship among soil biochemical properties, fungi and FUNGuild

The different environmental variables were significantly correlated with the variation of selected soil fungal genera (average relative abundance top 10) (Fig. 4A). The RDA demonstrated the relationship between soil properties and fungal communities, which had an eigenvalue of 0.2846 for the first axis and 0.0802 for the second axis, respectively. The vectors indicated the following: total N, total P, total K, pH, available N and soil temperature played a greater role than available K, available P and organic matter about sugarcane rhizosphere fungi genera. The soil fungus community were dominated by Basidiomycota and Ochrophyta, which showed stronger associations with higher pH, available N and soil temperature, while Chytridiemycota and Eukaryota were almost opposite to them, and then we used the Source Model of Plant Microbiome (SMPM) to estimate the proportion of sugarcane rhizosphere fungal communities from “adjacent period” and “unknown” sources and took the adjacent periods as the source and library in turn (Fig. 4B). The results showed that the related fungal communities mainly came from the transmission of adjacent periods, and the transfer proportion decreased gradually with the migration of time. In addition, Tremellales and Saccharomycetales, as the dominated order in fungi, were completely positive for associations with pH and available N, however, Sordariomycetes, Pseudallescheria and Colpodea showed stronger a negative association with pH and available N (Fig. 4D). To further explore the environmental driving factors of the sugarcane fungus community, we conducted a Mantal-Test analysis and analysed the correlation between the microbial matrix and the soil property matrix, and then we correlated distance-corrected dissimilarities of taxonomic and functional community composition with those of environmental factors. Overall, pH, TN and TK were the strongest correlates of taxonomic composition in the sugarcane rhizosphere soil (Fig. 4C), while no significant correlation was found for other soil factors (*p* < 0.05). In addition, soil temperature and pH were only weakly correlated to taxonomic and functional community composition. Besides, and except for available N, the correlations of others were not statistically significant.

**Fig. 4.**
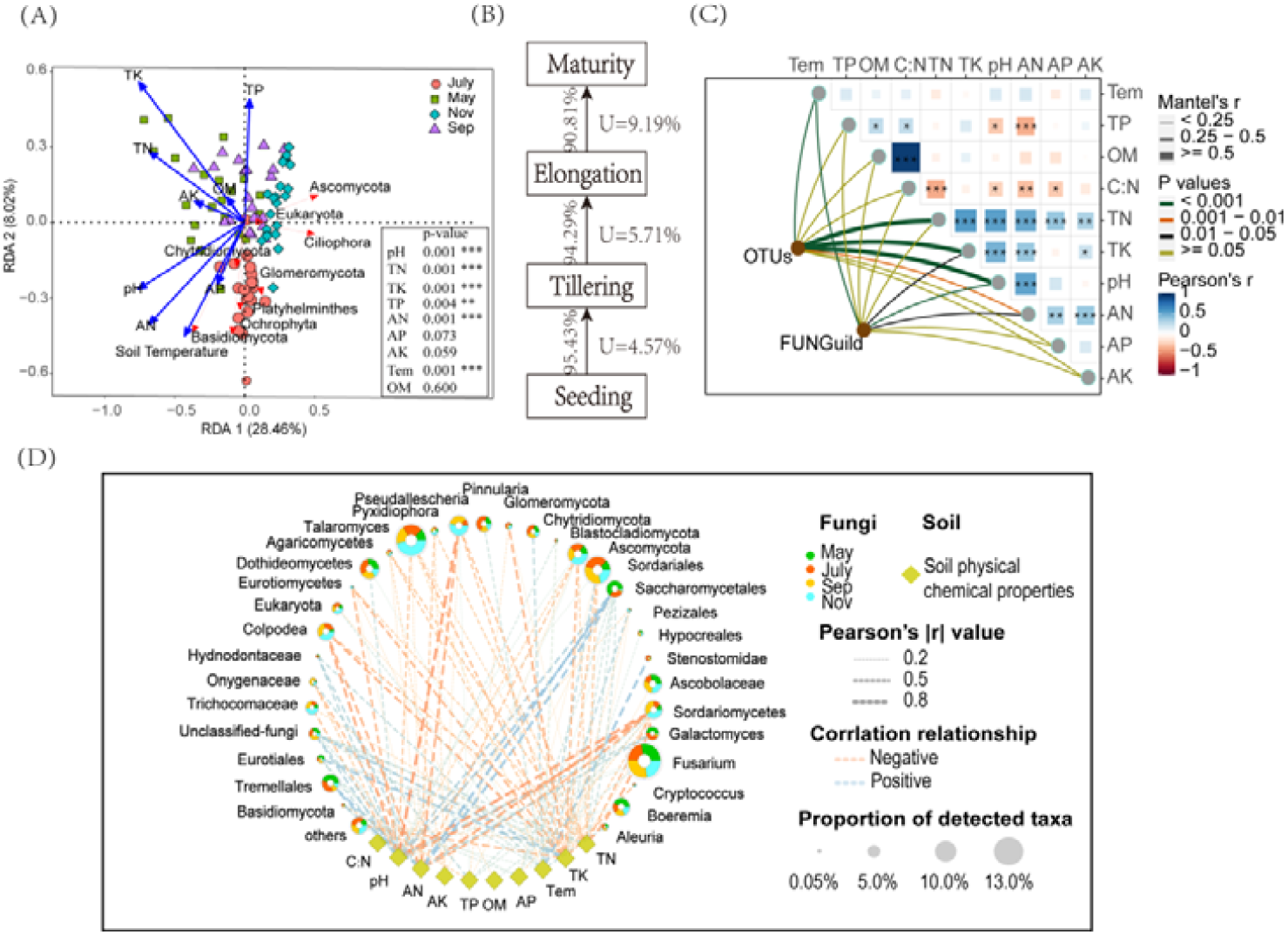
(A) Redudancy analysis (RDA) of soil properties and dominant Fungal phylum. In the part of bottom right, the soil properties were fitted to the ordination plots using a 999 permutations test (*p*-values), significance: *** 0.001, ** 0.01, * 0.05. (B) Source analysis of sugarcane rhizosphere microorganisms. source environment proportions for 84 samples of different seasons were estimated using Source Model of Plant Microbiome(SMPM). The number represented the proportion of fungal transmission in the adjacent period, and the letter U represented the proportion of microorganisms from unknown sources. (C). Environmental drivers of sugarcane rhizosphere fungal community composition. Pairwise comparisons of environmental factors are shown, with a color gradient denoting Pearson’s correlation coefficients. Edge width corresponds to the Mantel’s r statistic for the corresponding distance correlations, and edge color denotes the statistical significance based on 9,99 permutations. (D). Co-occurrence network of the fungus genus population (Relative abundance top 50) and soil properties in rhizosphere soils.

### Differences of sugarcane rhizosphere fungal communities in growth periods and co-occurrence network

We employed Manhattan plots to examine fungal community composition OTUs differences in sugarcane rhizosphere soil during the different periods (Fig. 5 and Table S1). The analysis revealed that Cilliopra was significantly higher in July. Furthermore, Basidiomycota, Ascomycota and Chytridiomycota enriched considerably in July compared to May periods (Fig. 5A). Additionally, Cilliopra and Ascomycota increased considerably in July relative to May, however Basidiomycota and Chytridiomycota decreased profoundly in May (Fig. 5B). In September, Basidiomycota and Ascomycota increased significantly, while Chytridiomycota diminished considerably in November (Fig. 5C). Meanwhile, Venn diagram analysis was adopted to have dipper understanding into the genera of fungal OTUs from one time point to another. The analysis demonstrated that between May and July, the highest amount of enriched fungal OTUs was detected, followed by September to November (Fig. 5E). However, between July and September the highest amount of depleted fungal OTUs were identified, followed by September to November (Fig. 5D). Meanwhile, the co-occurrence network showed the interactions of rhizosphere fungi in sugarcane during four critical periods, where the dominat fungus were belong to the mainly Talaromycetes *Talaromyces, Fusarium, Ascomycota, Sordariales* and *Pseudallescheria* (Fig. 5F). The role of each fungus in the network changed over time during different reproductive periods, with the degree and importance changed over time. Among the four critical fertility networks, May has the largest average density and mean degree, followed by November (Table S9).

**Fig. 5.**
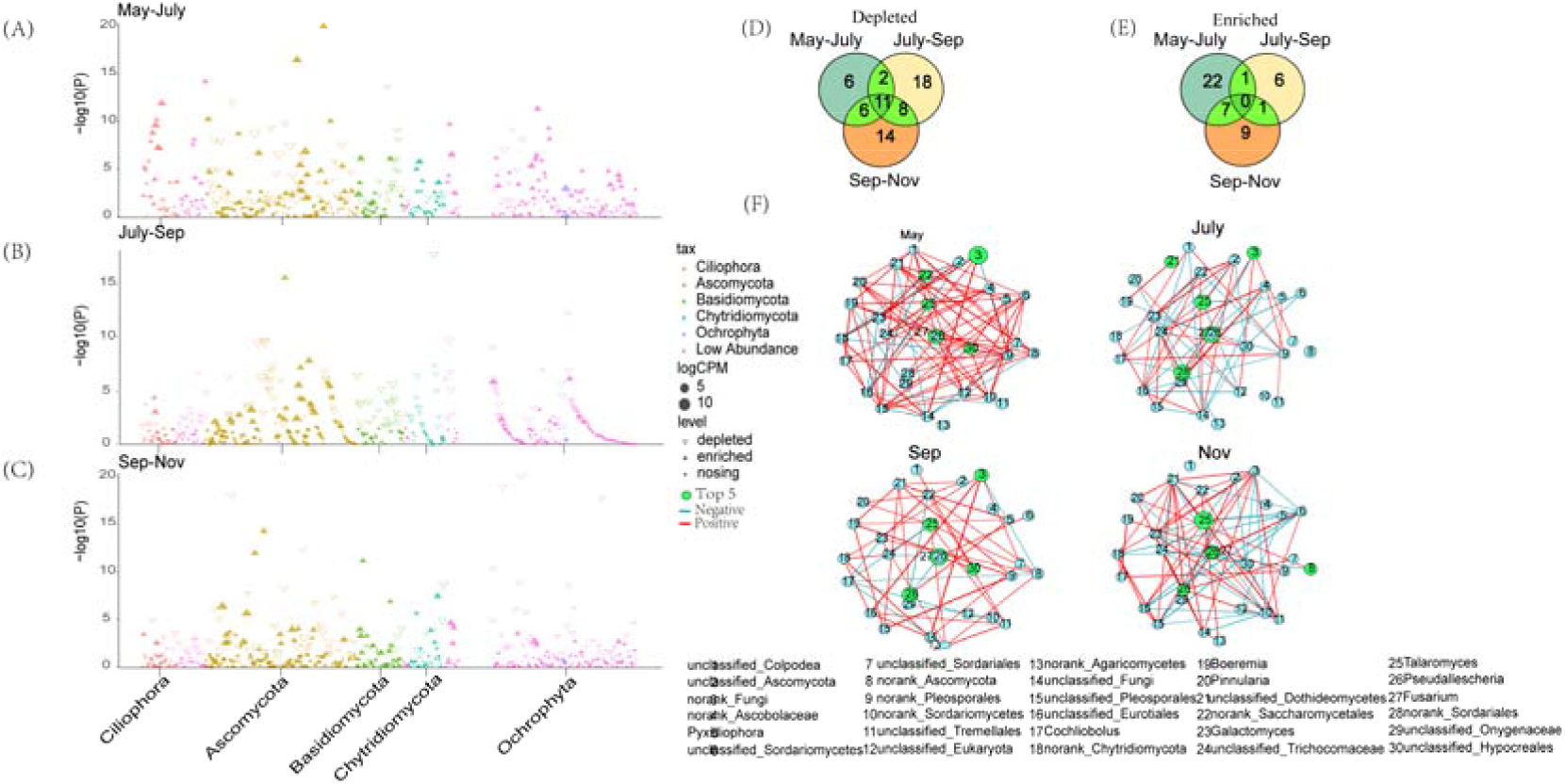
Taxonomic of differential fungi between the two periods root microbiota. Manhattan plot showing OTUs enriched or decreased in sugarcane in different stages. Each dot or triangle represents a single OTU. OTUs enriched or decreased in sugarcane rhizosphere are represented by filled or empty triangles, respectively (FDR adjusted *p* < 0.05, Wilcoxon rank sum test). OTUs are arranged in taxonomic order and colored according to the phylum. (A) May vs. July; (B) July vs.September; (C) September vs. November. (D) Venn diagram of fungi decreasing between successive stages at the genus level. (E) Venn diagram of fungi enriched between successive stages at the genus level. (F) Co-occurrence network based on spearman correlation (Top 30 fungal genera, |r| > 0.4, *p* < 0.05), the size of the circles represents the relative abundance of the fungal genera and the thickness of the lines represents the size of the correlation.

### Construction of structural equation models related to pokkah boeng disease of sugarcane

We constructed a SEM to assess the direct and indirect effects of soil properties (pH, AN and TN), microbial genera (*Fusarium* and *Talaromyces*) and microbial diversity (ACE and Shannon) on the incidence of sugarcane (Fig. 6, Fig. S5 and Table S2). We used a multigroup modelling method to assess which relationships between soil properties, microbial communities, and sugarcane disease during grow process. The results of model-1 showed pH significantly and directly affected the abundance of *Fusarium*, sugarcane disease and fungal diversity, however, the difference was that pH negatively regulates the disease rate of sugarcane (the lower the pH, the more disease occurs)(Fig. 6A and Table S3). Meanwhile, AN, TN and disease had similar significant regulatory relationships as pH and disease (Fig. S5B and S5C, Table S6 and S7,). In addition, soil temperature had effects on the diversity of rhizosphere fungi, and at the same time had both direct and indirect effects on the abundance of *Fusarium* and *Talaromyces*.

**Fig. 6.**
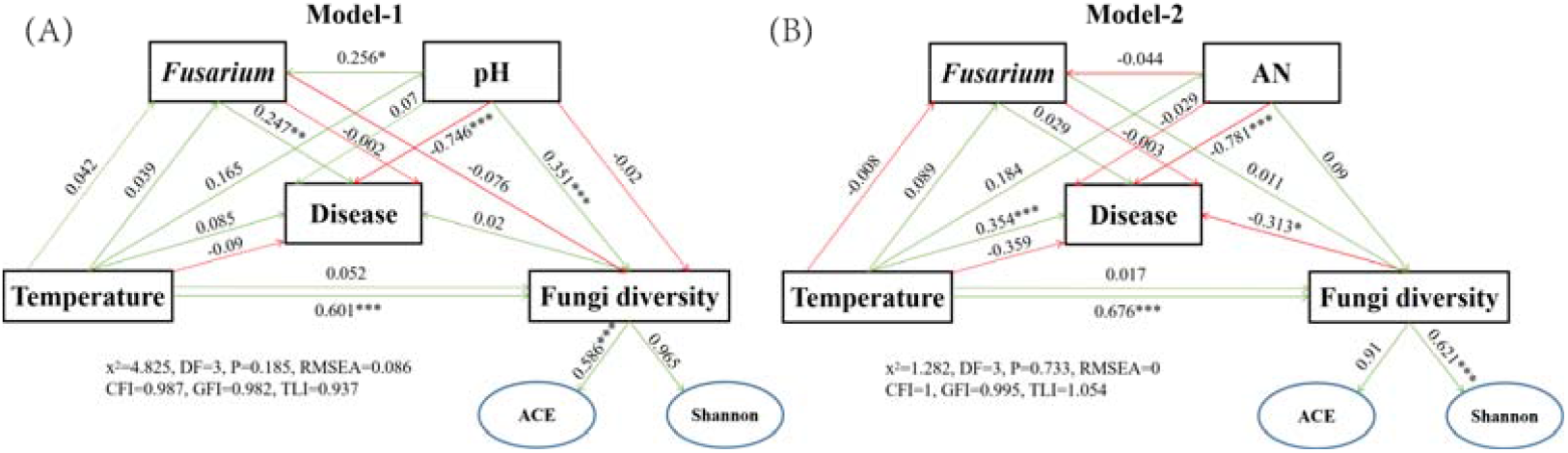
Structural Equation Modeling (SEM) analysis showed the contributions of *Fusarium*, temperature (soil temperature), soil and fungi diversity to the sugarcane disease. Disease was pokkah boeng disease of sugarcane. Soil was represented by mainly compositions including pH and AN (Available Nitrogen); Fungi diversity included ACE and Shannon index. The number above the arrow indicates the normalized path coefficient. *p-*value: * < 0.05; ** < 0.01; *** < 0.001.

### Assembly process of sugarcane rhizosphere fungi community

The results showed that the mean nearest-taxon-index (NTI) was greater than 0, and the mean nearest taxon distance between samples (β-NTI) values among the samples from the critical fertility period of sugarcane were all mainly concentrated in the interval of -2 to 2, indicating that the changes in microbial community structure during the critical fertility period were mainly influenced by stochastic factors. The existence of a certain number of samples with β-NTI values greater than 2 among the soils in different periods indicated that the changes in rhizosphere microbial community structure of some soil samples was mainly influenced by deterministic factors (Fig. 7B). In addition, the β-NTI values of samples between months (May-July, July-September and September-November) were between (−2, 2), indicating that the changes in the rhizosphere microbial community structure of sugarcane with time were mainly influenced by stochastic factors, but there were differences in the contribution of stochastic factors with time (Table S8).

**Fig. 7.**
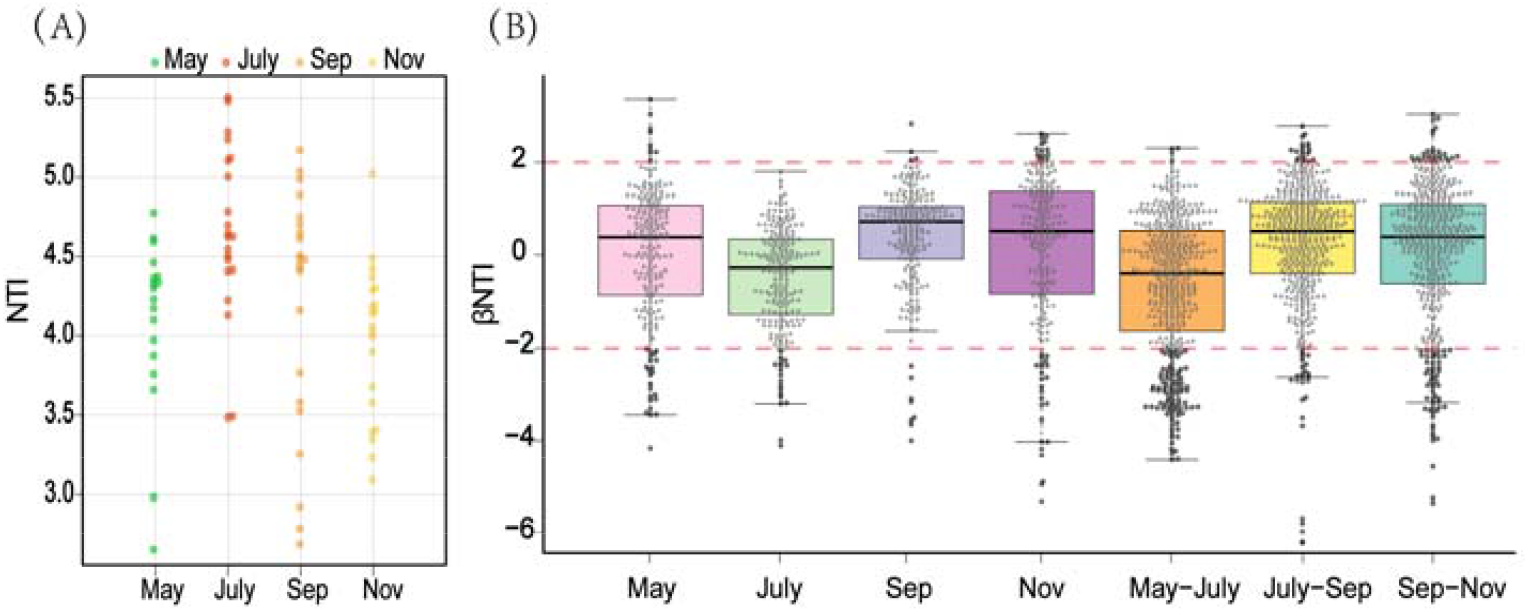
The distribution of NTI values (A) and box plots of β-NTI (B) for rhizosphere soil samples of sugarcane at different fertility stages, with different colors representing sample groups at different periods and red dashed lines marking the intervals of -2 and 2.

### Functional and evolutionary relationships of the major fungal genera of rhizosphere soil

As for functional classification, the fungal communities in rhizosphere soil were classified by using the trophic mode (Fig. 8). To further visualised the relationship among crucial fungal communities, a Maximum Likelihood phylogenetic tree was conducted. Following the procedure, the top 30 genus, were categorised into five guilds. They were divided into Undefined Saprotroph, Dung Saprotroph, Animal Pathogen, Plant Pathogen, Endophyte, respectively. Based on the classification, the results revealed that 33.3% of these genus belonged to Undefined Saprotroph, 3.3 % belonged to Plant Pathogen, 6.6 % belonged to Animal Pathogen, 6% belonged to Dung Saprotroph, 3.3% belonged to Endophyte while 46.6% of the top 30 genus were unclassified. Among the top 30 genera, they come from 8 different phyla, and the five that had been identified were Ascomycota, Chytridiomycota, Ochrophyta, Ciliophora, Basidiomycota, respectively. The plant pathogenic fungi observed in this study were Cochliobolus, and it was worth noting that the abundance of *Fusarium* genera was higher dominant in July and November. In addition, according to the results of the FUNGuild classification difference (Fig. S6), it was found that in the comparison of the May and July seasons of sugarcane growth (Fig. S6A), there were more classifications with significant differences (*p* <0.05), they were Dung Saprotroph-Soil Saprotroph-Wood Saprotrop, Animal Pathogen-Endophyte-Lichen Parasite-Plant, Orchid Mycorrhizal-Plant Pathogen-Wood Saprotroph, Endophyte-Plant Pathogen, Animal Pathogen-Soil Saprotroph respectively. However, with time, the FUNGuild classification with significant differences gradually decreased and the FUNGuild classification with significant differences but not clear appeared (Fig. S6B and S6C).

**Fig. 8.**
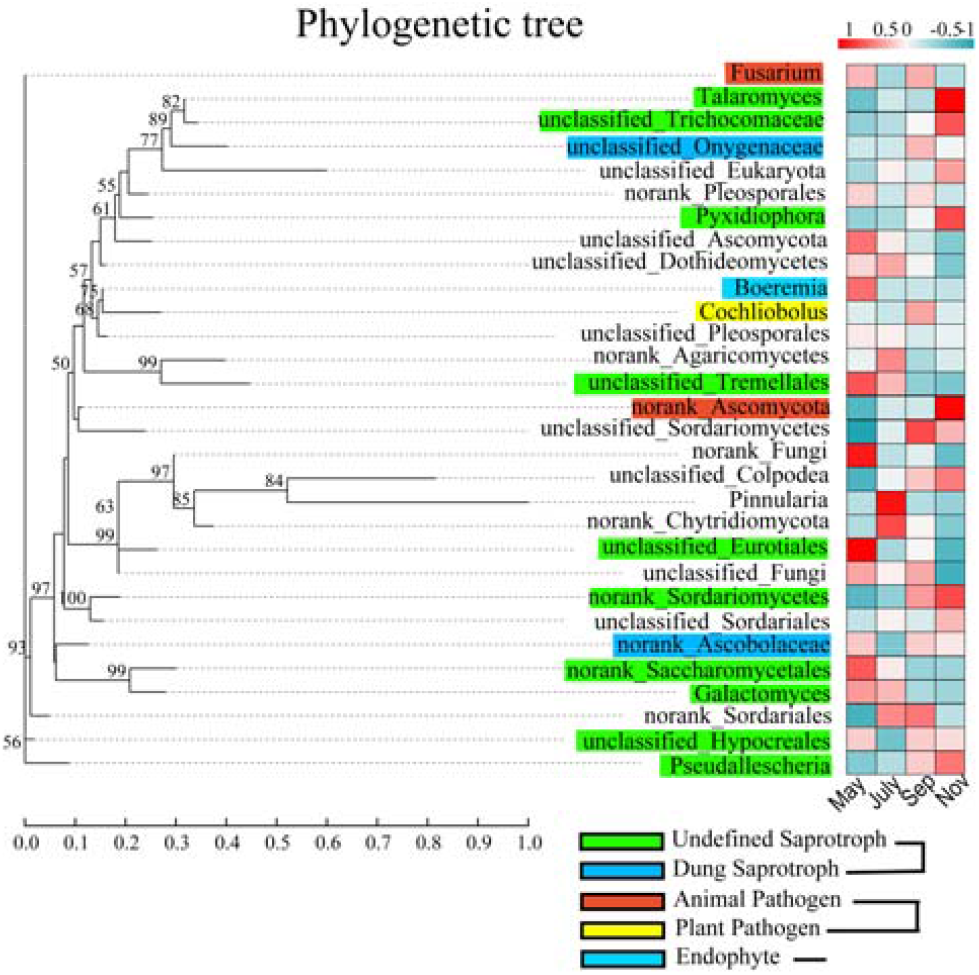
Phylogram with fungal guilds of the top 30 fungal genera,color the branches according to the phyla level to which the species belongs; Maximum Likelihood tree for the sequences obtained through high-throughput sequencing. The relative abundance data were normalized using the Z-score method, and then the average of the relative abundance of each genus for each group of samples was plotted as a heat map, the bar chart showed the percentage of Reads for the species in different stages. Guild annotation used FUNGuild database.

## DISCUSSION

Soil microbial communities associated with plants can have strong influences on plant growth as well as contribute to soil health and sustainable production (48-50). Therefore, understanding the temporal progressions of the root microbiota is an important aspect for plants and soil environmental improvement (51,52). Previous researches showed that plants root microbiota composition varied with plants developmental stage (53,54), but these works were carried out either with other model crops or under greenhouse conditions. Our results provide a specific characterisation for the rhizosphere microbiota of the sugarcane whole growth period in the field and reveal views into how sugarcane growth and soil environment affect the development of the rhizosphere microbiota. We show temporal shifts in fungal community composition during the life of sugarcane, and these results are missing in related research on sugarcane in the field (Fig. 1A and 1B). The results show that the nutrients of sugarcane rhizosphere soil have significant changes in different growth periods (*p* < 0.05), which might be due to the difference in sugarcane requirements for different nutrients at different growth stages (55). Moreover, although the alpha diversity of the sugarcane rhizosphere fungal community varies significantly between different growth periods, it has a gradually stable trend (Fig. S1A-S1D). In addition, the species composition show that rhizosphere fungi were mainly Ascomycota and Basidiomycota during the growth period of sugarcane (Fig. 3A), which was similar with the research data of Zeng et al., (2020) (56). Stursova et al., (2012) also found that compared with Basidiomycota, Ascomycota are more involved in cellulose decomposition (57). Meanwhile, Chytridiomycota, Eukaryota, Glomeromycota and Ochrophyta which had a lower relative abundance also showed regular and significant changes over time (*p* < 0.05). Determining the role, if any, of these low-abundance microbes in responding to changes in plant diseases, soil health, and so on will be a fascinating challenge for future studies. Venn diagram shows that 511 OTUs shared by sugarcane during the growth periods (Fig. 3B), the number of OTU unique to each of these periods varies considerably (May: 25, July: 53, Sep: 20, Nov: 15). To further explore, we used the source model of microorganisms to analyse the fungal transmission ratio between seasons (Fig. 4B). The results imply that 90% of the fungal population is passed to the next period in each season. This indicates that after the formation of the sugarcane rhizosphere fungal flora, although different microorganisms would be recruited or consumed at different periods, the overall structure would remain stable. This result also verifies the idea that plants can maintain resident soil fungal populations, but not non-resident soil fungal populations, as has been previously verified in the model plants (5). Besides, there is a strong correlation between soil environmental factors and sugarcane rhizosphere fungi populations in different seasons, RDA and network map show that TN, TK, PH and AN are the main factors driving sugarcane fungal communities (Fig. 4A and 4D). Those soil nutrients directly or indirectly affected the survival and growth of sugarcane fungal communities. This is similar to the results reported by Zhang et al., (2016) They indicated that organic matter, total N and total P significantly affected soil fungal community composition in the southeastern Tengger Desert (58). To further explored the influence of these nutrient factors on the fungal community during the growth period of sugarcane, we use the mantal-test to calculate the correlation between three matrices and found that TN, TK and pH were still the main environmental factors driving the OTU composition of the fungal community (Fig. 4C). There are more genus that were decreasing in relative abundance in the rhizosphere over the life cycle of sugarcane, while fewer phyla were increasing in relative abundance. It indicates that a smaller proportion of the soil fungi were enriched by the sugarcane, while more fungus taxa were depleted (Fig. 5D and 5E). This is similar to the results of tracking changes in the rice root microbial flora (51). These data reinforce the separation role that the growth period to distinguish plant rhizosphere microbial flora as observed in other studies (51), but it needs more research to illustrate. During the whole growth period of sugarcane, there are 11 fungal genera that had been decreasing, mainly including Ascomycota, Chytridiomycota, Eurotiales, Hypocreales, Sordariomycetes, Tremellales. We speculate that the decline of these fungal genera is closely related to the genotype and developmental stage of sugarcane (62, 63). This requires us to conduct further verification. Such results provides a list of sugarcane rhizosphere fungi which should now be targeted to elucidate their potential functions in the roots of sugarcane (symptom-less colonizers vs. plant growth-promoters vs. pathogens) by targeted separation and sequencing technology. Furthermore, according to our results, Pseudallescheria had been enriched during the first three growth periods of sugarcane growth (May-Sep). However, *Nectriaceae* is significantly (*p* < 0.05) enriched in the tillering (July), elongation (Sep) and maturation (Nov) stages (Fig. 5A-5C). *Pseudallescheria* is a filamentous pathogenic fungi, *Pseudallescheria* is not only a potential human and animal pathogen, but also exists in the soil environment (64,65). It has special public health importance because it causes localised as well as disseminated infections in both immunocompromised and immunocompetent hosts (66). Studying the time trend of this pathogenic fungus in the sugarcane rhizosphere has a great significance for the defense of certain new diseases that occur in the early growth of sugarcane in the future. Similarly, the ascomycete family *Nectriaceae* also includes numerous important plant and human pathogens (67). The enrichment of *Nectriaceae* may be potentially related to certain diseases that occurred in the late stage of sugarcane growth. However, evidence of pathogenicity does not necessarily exist, which needs to be established through more in depth and mechanistic studies. Nonetheless, our findings reveal the most basic information and provide the possibility of sugarcane disease research. In addition, it is found in the co-occurrence network that some fungal interactions between genera disappeared and then reappeared with time, presumably because the “autonomous consciousness” of sugarcane and environment factors adjusted the balance of fungal interactions. Whether this phenomenon is associated with the development of pokkah boeng disease remains to be proven.

Structural equation models (SEMs) indicates that soil temperature is an important factor affecting the α-diversity of rhizosphere fungal communities of sugarcane, which show significance in all models that we construct (Fig. 6 and Fig. S5). Soil temperature changes brought about by seasonal changes had a certain perturbing effect on the rhizosphere community (68). In addition, the strong direct negative effects of soil pH, AN and TN with the disease status of sugarcane maybe suggest that soil infertile would increase the likelihood of sugarcane disease (69). At the same time, *Fusarium* and *Talaromyces*. also respond to changes in the soil nutrient environment, thus maybe interacting adversely with the plant. The high quality genome sequence of *Fusarium andiyazi* in China published by Bao et al., (2021) provides more clarity on the role of *Fusarium* in sugarcane pokkah boeng disease (70). In addition, the community assembly process of rhizosphere fungi, the results show that the majority of the βNTI values among the samples is less than |2| (Fig. 7B), indicating that the changes produced by the rhizosphere fungal community of sugarcane over time were mainly influenced by stochastic factors (71), but the stochastic contribution increase and then decrease with the growth and development of sugarcane, and finally reached the lowest value at the maturity stage (Table. S8). This maybe due to stochastic changes in the probability distribution and the relative abundance of species of rhizosphere fungi in the early stages of sugarcane growth (ecological drift) (72), and the formation of an adaptive system of root and soil environment in the later stages of sugarcane growth, resulting in abiotic and biotic factors playing an increasingly important role in the presence or absence and relative abundance of rhizosphere fungi. The construction of phylogenetic trees and the application of FUNGuild analysis in the study of rhizosphere fungal communities provides a way to simplify and visualize complex communities (Fig. 8 and Fig. S6). FUNGuild analysis reveals that the abundance of fungi showed differential responses across seasons, for example *Fusarium* is abundant in May and September and *Talaromyces* in November. these result correlate with the work of Li et al., (2021) who reported that some potential biocontrol genera change plus with plant growth and tillage (73). While showing dominant species richness, it also provided new ideas for seasonal prediction of rhizosphere pathogens of sugarcane and disease control during the different growth period, such as annotated phytopathogens (*Cochliobolus*) that may directly or indirectly contribute to the disease of sugarcane in a given season.

## CONCLUSION

When focusing on soil productivity such as plant yield, rhizosphere microbiomes are also an important factor and closely related to plant health. In this field study, both sugarcane growth period and soil traits strongly influenced the composition and function of rhizosphere fungal communities. The pH, TN, TK and AN decreased significantly during the growth of sugarcane, the dominant fungal phyla were Ascomycota, Basidiomycota and Chytridiomycota, and these variations could be partially explained by plant rhizosphere dynamics. Using SEM, we found that diseases showed significant and strong negative effects with selected soil traits, soil temperature had a strong and direct positive effect on fungal α-diversity, while *Fusarium* and *Talaromyces* also had some direct and indirect associations with sugarcane diseases indicating that changes in soil nutrients affect plant health and rhizosphere fungal interactions. The variation in rhizosphere fungal structure was mainly influenced by stochastic factors, and the variation in stochastic contribution may be due to the coupling of sensitive fungal genera or modules in the sugarcane rhizosphere fungal network with sugarcane growth dynamics to coordinate the growth process of sugarcane. These results contribute to the understanding of plant-rhizosphere fungal interactions and provide additional opportunities to develop greener and more productive agricultural practices.

## AUTHOR CONTRIBUTION STATEMENT

All authors contributed to the intellectual input and provided assistance to this study and manuscript preparation; Z.Y. and Z.P. designed the research and conducted the experiments; Q.L. analyzed the data and wrote the manuscript; Y.L., Fallah.N., W.L. and C.H. reviewed the manuscript; Z.Y supervised the work and approved the manuscript for publication.

## ACKNOWLEDGMENTS

Thanks for the data analysis provided by the free online platform of the Magi Cloud platform (www.majorbio.com).

## SUPPLEMENTARY MATERIAL

Fig. S1 | Box plots of rhizosphere microbial α-diversity of sugarcane in different seasons.

Fig. S2 | The rarefection curve of fungal alpha diversity index.

Fig. S3 | Linear relationships among selected microbial characteristics and soil physicochemical variables (*p* < 0.05).

Fig. S4 | The box plot shows the changes of rhizosphere fungi in four different sugarcane growth stages.

Fig. S5 | Structural Equation Modeling (SEM) analysis showed the contributions of *Talaromyces* (A and B), *Fusarium* (C), temperature, soil and fungi diversity to the sugarcane disease.

Fig. S6 | The functional classification with significant functional difference phases in different seasons was obtained based on FUNGuild database.

Table S1 | Summary table of fungal genera declining or enriched in sugarcane roots in the comparison between seasons.

Table S2 | Model parameters summary table

Table S3 | Summary of model-1 internal path coefficients and normalized coefficients.

Table S4 | Summary of model-2 internal path coefficients and normalized coefficients.

Table S5 | Summary of model-3 internal path coefficients and normalized coefficients.

Table S6 | Summary of model-4 internal path coefficients and normalized coefficients.

Table S7 | Summary of model-5 internal path coefficients and normalized coefficients.

Table S8 | Summary of mean NTI and βNTI during fungal assembly.

Table S9 | Network structure parameters summary table.

## DATA AVAILABILITY S TATEMENT

The complete data sets generated in our study have been deposited in the NCBI Sequence Read Archive database under BioProject ID PRJNA721464.

